# Organoid generation from mouse mammary tumors captures the genetic heterogeneity of clinically relevant copy number alterations

**DOI:** 10.1101/2023.01.29.526141

**Authors:** Katherine E. Lake, Megan M. Colonetta, Clayton A. Smith, Kenneth Martinez-Algarin, Kaitlyn Saunders, Sakshi Mohta, Jacob Pena, Heather L. McArthur, Sangeetha M. Reddy, Evanthia T. Roussos-Torres, Elizabeth H. Chen, Isaac S. Chan

## Abstract

Breast cancer metastases exhibit many different genetic alterations, including copy number amplifications. Using publicly available datasets, we identify copy number amplifications in metastatic breast tumor samples and using our organoid-based metastasis assays, and we validate FGFR1 is amplified in collectively migrating organoids. Because the heterogeneity of breast tumors is increasingly becoming relevant to clinical practice, we demonstrate our organoid method captures genetic heterogeneity of individual tumors.

## Main Text

While breast cancer is the most prevalent cancer among women, most patients are diagnosed with early-stage breast cancer and cured by multi-modality treatment [1]. However, around 10% of patients will develop metastatic breast cancer (MBC), which is the main driver of breast cancer related deaths [2, 3]. A reason for therapeutic resistance of MBC is partly due to the relative lack of targetable genetic vulnerabilities that act as intrinsic mediators of breast cancer cell metastasis. Recent literature suggests that cancer cell metastasis is defined by copy number alterations and not sufficiently by genetic mutations alone [4–7].

To identify potential copy number amplifications involved in metastatic breast cancer, we used the Project GENIE database [8] and cBioPortal [9] for analysis, genes with copy number alterations in invasive ductal carcinoma (IDC) patient samples were selected. Genes were filtered first by copy number amplification in >10% of metastatic IDC samples, defined as having distant organ metastasis, unspecified metastasis site, or lymph node metastasis. From those, the top 10 differentially amplified genes in metastatic IDC over primary IDC samples were identified. In order by logarithmic ratio, they include ADGRA2, RAD21, PAK1, FGF4, NSD3, FGF19, FGF3, CCND1, FGFR1, and MYC (**Fig. 1a**).

**Figure 1:**
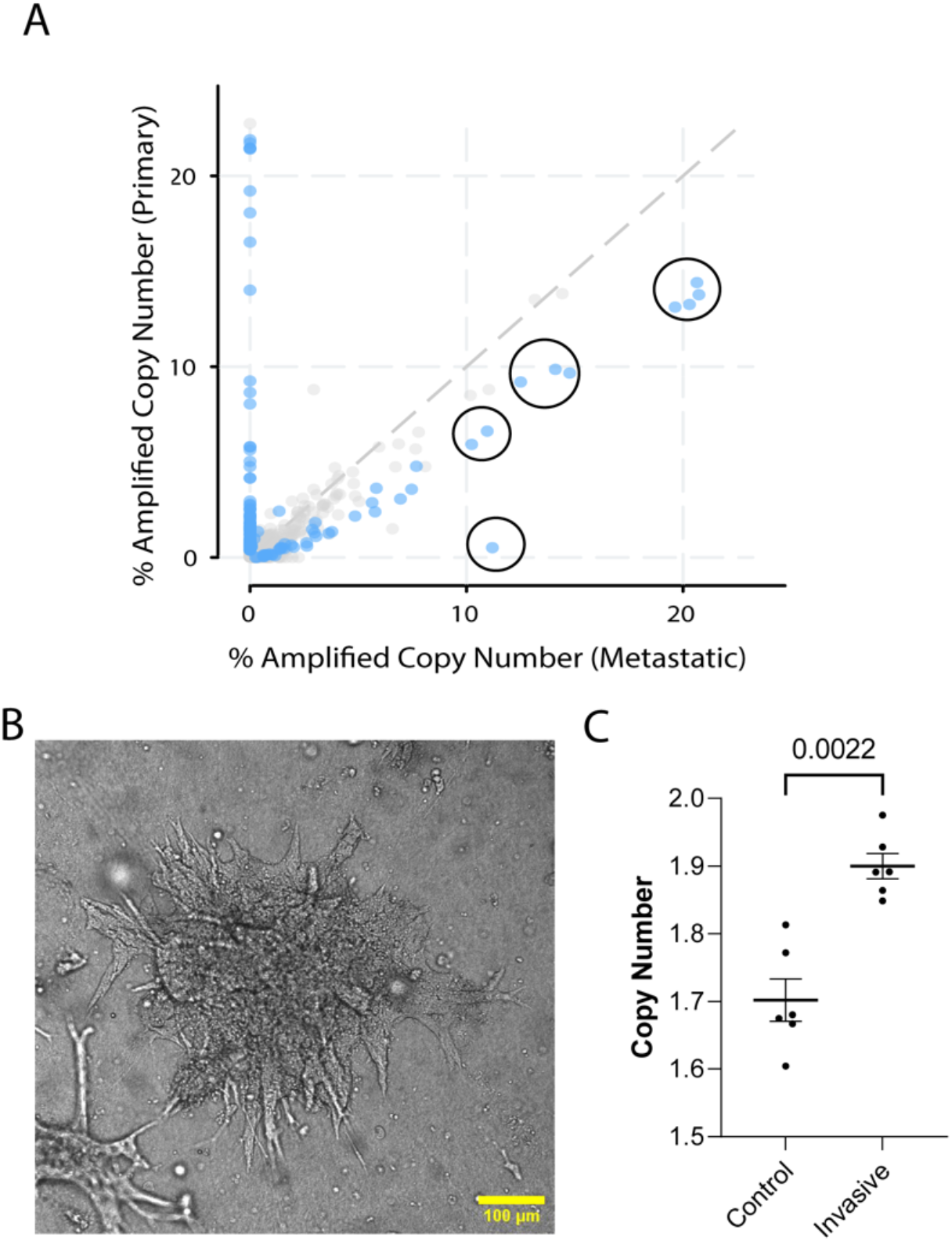
FGFR1 is copy number amplified in both human metastasis and in-vitro model of metastasis. **A)** Copy number amplified genes in IDC patients found in distant organ metastasis, unspecified metastasis site, or lymph node metastasis compared to primary breast cancer samples in the GENIE database. Of the genes amplified in greater than 10% of metastatic samples, the top 10 differentially amplified in metastatic samples are highlighted. These include ADGRA2, RAD21, PAK1, FGF4, NSD3, FGF19, FGF3, CCND1, FGFR1, and MYC. **B)** Mouse PyMT organoid invading in type 1 collagen matrix. **C)** FGFR1 copy number is amplified in invading organoids vs. control organoids from 2 PyMT mouse primary mammary tumor samples (Mann-Whitney, p=0.0022).

Of the genes found in Figure 1a, FGFR1 is the most clinically mature target, with multiple therapeutic inhibitors either in clinical trials or in the drug development pipeline [10]. We reasoned if FGFR1 copy number is amplified in metastatic lesions, it could potentially be amplified starting at the earliest stages of metastasis, invasion out of the primary tumor. To test whether FGFR1 copy number amplification is associated with invasion, we generated mammary tumor organoids derived from MMTV-PyMT mice using differential centrifugation [11–13]. Organoids were then randomly split into a control group which were grown in suspension and an experimental group grown in 3D culture. Growing organoids in 3D culture allows ex vivo assessment of their invasive potential through collective migration into collagen [11–13]. In brief, the organoids were either kept in media suspension or embedded in 3D collagen I gels and assessed for invasion (**Fig. 1b**). Both the control and experimental groups were allowed to grow for 48hrs. Then, we isolated genomic DNA from both groups. Interestingly, invasive organoids retain their morphology after removal from collagen gels (**Supplementary Fig. 1**). We then determined gene copy number of FGFR1 in control and invasive samples using ddPCR. We found that invasive organoids have statistically significant copy number amplification compared to control organoids (Mann-Whitney, p=0.0022) (**Fig. 1c**). These findings demonstrate that higher FGFR1 copy number amplification correlates with organoid invasion, suggesting that FGFR1 is involved in the earliest stages of metastasis in addition to the developed metastases. However, although we observed increased copy number of FGFR1 in invasive organoids over non-invasive organoids, the average copy number of each experimental condition was below 2, indicating heterogeneity of FGFR1 amplification between organoids in the sample.

Recent and early literature have suggested that metastatic events are spurred by only a small number of cells from genetically heterogeneous primary tumors, and that only a few genetically predisposed cells are capable of metastasis [14, 15]. However, the functional value of heterogeneity within tumors has been hard to model [16, 17] To better understand the genetic heterogeneity of our organoid generation method, we processed mouse mammary tumors into tissue, pooled organoid, single cell digest, and single organoid samples (**Fig. 2a**). Of the top 10 differentially amplified genes in metastatic over primary breast cancer, as identified in Figure 1a, we focused on four genes, ADGRA2, FGFR1, NSD3, and PAK1, which have been reported to reside in chromosomal regions copy number altered in MMTV-PyMT mouse tumors [18]. Mammary tumor samples were digested to isolate gDNA and identify the copy number of ADGRA2, FGFR1, NSD3, and PAK1 using ddPCR (**Fig. 2b**, **Supplementary Figs. 2-4**). Notably, in all but PAK1, tissue copy number varied significantly from pooled organoid samples (Mann-Whitney, p<0.05). Pooled organoid samples include cancerous mammary epithelium and exclude any stroma, muscle, or other cells that may be included in tissue samples [11, 13]. Thus, copy number reads from pooled organoid samples provide a more accurate representation of cancer epithelial cells within a tumor instead of a mixture of epithelial, stromal and immune cells. Unsurprisingly, no single cell digest copy number was statistically distinct from the pooled organoid sample from the same tumor in any tested gene. However, single cell digests lose the potential for functional testing and risk skewed genetic profiles through imperfect digestion or straining.

**Figure 2:**
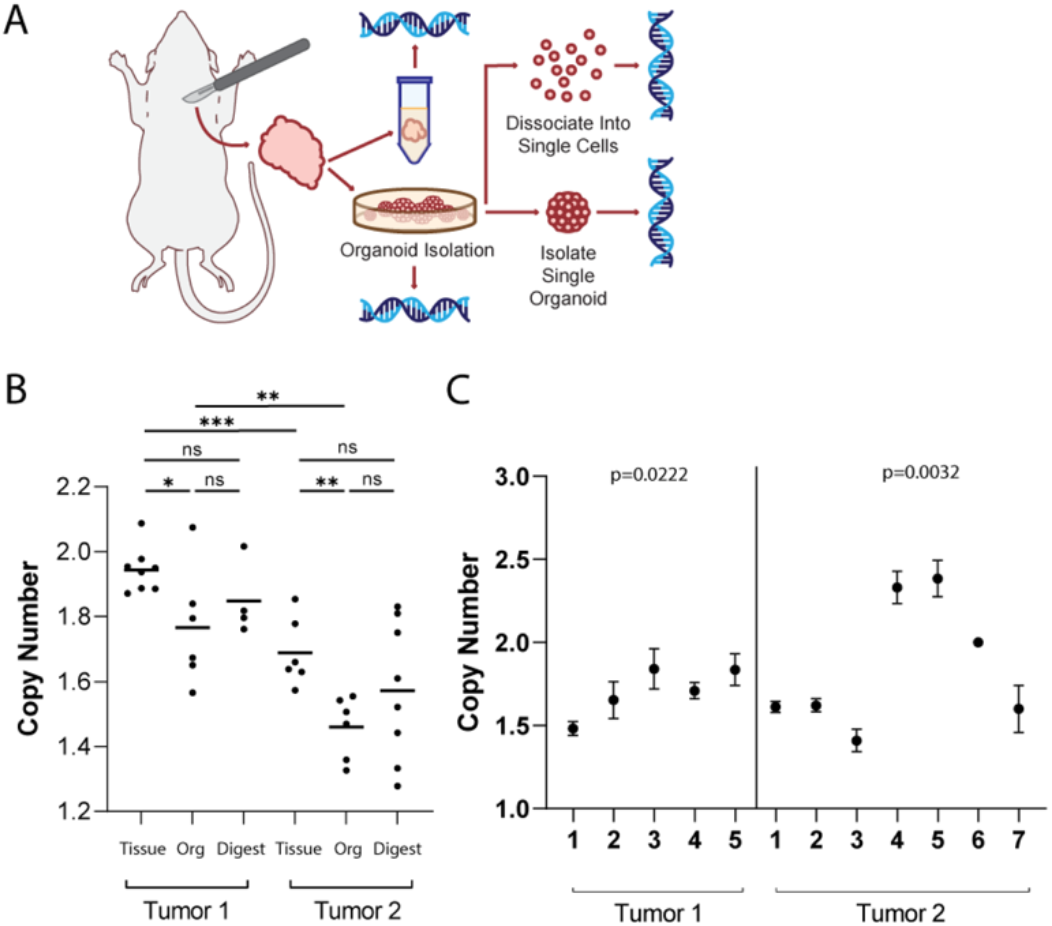
Organoid generation retains clinically relevant tumor genomic heterogeneity. **A)** Schema of workflow for sample generation. Large primary mammary tumors are dissected from the fat pad. Tissue segments are excised from 4 distinct regions of the tumor to ensure adequate sampling. Organoids generated from the tumors were sampled and dissociated and strained to single cells, taken as pooled organoid samples, or isolated to single organoids. All samples were then used to isolate gDNA and perform ddPCR. **B)** FGFR1 copy numbers in tissue, pooled organoids (Org), and single cell digests (Digest) from two different mice. Tissue sample copy number alterations are statistically different than pooled organoid samples (Mann-Whitney, p<0.05). **C)** FGFR1 copy numbers in single organoids from tumors from two different mice. Among each tumor there exists statistically distinct copy numbers (Kruskall-Wallis, p=0.0222; p=0.0032).

Interestingly, pooled organoid samples differed from a normal diploid state, suggesting the sample was heterogeneous and included cells with copy number variations reflective of an aneuploid state (ie. CNV 0 or 1 or 3). To identify whether this heterogeneity was consistent at the organoid level, we isolated gDNA from single organoids and tested the copy number of ADGRA2, FGFR1, NSD3, and PAK1 **(Fig. 2c, Supplementary Figs 2-4).** Copy number of select genes differed widely among individual organoids. Additionally, copy number was rarely identified as exactly 2 in single organoids, suggesting that not only are organoids genetically distinct from each other, but they also must contain cells with copy number alterations in different proportions. In summary, our organoid isolation method captures both the intra- and inter-organoid genetic heterogeneity in tumor samples.

## Discussion

In this study, we determine ten copy number amplifications enriched in metastatic site tumors over primary breast tumors using Project GENIE [8]. In assessing FGFR1, we found that it is amplified in invading organoids, suggesting the importance of FGFR1 in the early stages of metastasis. Lastly, given the increasing clinical importance of cancer epithelial heterogeneity within the breast tumor, we demonstrate that this method of organoid generation captures intratumoral genetic diversity between individual organoids. As far as we know, this is the first study to analyze the copy number state of individual breast organoids. Further, one main challenge to using organoids in preclinical models is genetic drift [19]. Here we show that isolating organoids without passaging captures both intratumoral heterogeneity and reduces the chance of genetic drift as organoids are generated and used soon after tumor digestion. Given that functional models are lacking in the literature to test the impact of tumor heterogeneity on tumor metastasis [20] and immune interactions [21], this model could be useful for further experimentation.

This work is limited by the number of tumors used and the number of genes experimentally validated. Future work should focus on additional functional validation of the role of FGFR1 in metastatic development. The implications of FGFR1 knockout should be assessed in both in-vitro tumor organoid invasion models as well as in-vivo metastasis models to better understand the necessity and sufficiency of FGFR1 in the metastatic cascade. In addition, only mouse models were used to assess copy number heterogeneity, which may differ from human tumors. Further, single organoids are typically comprised of 50-100 cells, and thus the quantity of gDNA extracted is limited, restricting the replicate number, positive droplets in ddPCR, and ultimately statistical power of the analysis. Overall, our work contributes to the growing need for improved modeling of intratumoral cancer epithelial cell heterogeneity, which has broad implications on clinical practice and cancer biology.

## Methods

### Copy Number Amplifications in Metastatic over Primary Breast Cancers

We analyzed the GENIE dataset from invasive ductal carcinomas (IDC) patients. We obtained copy-number amplification (CNA) information from metastatic samples defined as distant organ metastasis, unspecified metastasis site, or lymph node metastasis and primary breast tumors. The alteration frequency was analyzed by using cBioPortal.

### Animals, Tumor, Organoid, and Cell Samples

11–14-week-old MMTV-PyMT mice with large (>0.5cm) palpable mammary tumors were identified. Mice were sacrificed according to IUCAC guidelines with CO2 asphyxiation and secondary cervical dislocation. Tumors were dissected and organoids generated following the protocol described previously [11]. For Figure 2a, tumors were dissected into quadrants, and samples taken as described in schema. Single cells were strained and harvested after Tryple™ Express (Gibco™; cat: 12605036) digestion and visual verification of single cell dissociation. Organoid invasion assay was performed as described previously [11]. Collagenase was used to digest 3D collagen I gels to isolate invasive organoids. Single organoids were isolated using P20 pipettes set to ~5uL until a single organoid was isolated into a well and visualized via microscope. If organoid density was too high for single organoid isolation, ~20uL was diluted into 500uL PBS in a 12-w plate. Microscope verification was performed for each single organoid. Genomic DNA was isolated using Quick-DNA Miniprep kit (Zymo Research).

### Droplet Digital PCR

Primers and probes for ddPCR for reference gene (RPP30) and target genes (ADGRA2, FGFR1, NSD3, PAK1) were purchased from Bio-Rad Laboratories (Assay IDs: dMmuCNS822293939, dMmuCNS263266645, dMmuCNS890129559, dMmuCNS681547140, dMmuCNS429051281, respectively). ddPCR supermix (no dUTP) (Bio-Rad; cat: 1863024), HaeIII restriction enzyme and rCutsmart buffer (NEB; cat: R0108S), and nuclease-free water were mixed with primer/probe for target and reference gene according to manufacturer recommendations. Droplets were generated using the QX200 droplet generator (Bio-Rad) and subsequently thermocycled according to manufacturer recommendations. Following PCR amplification, CNA data was acquired using the QX200 droplet reader (Bio-Rad) and analyzed with QuantaSoft software (BioRad). Droplets were plotted based on their fluorescence amplitude of each probe, high in positive droplets and low in negative droplets. Thresholds to determine positive and negative droplets were visually set between the two clusters. Technical duplicates were performed for every sample, and those with copy number reads greater than 20% apart were excluded from analysis. Unpaired non-parametric t-tests (Mann-Whitney tests) were performed for each comparison of copy numbers between conditions. Kruskall-Wallis tests were performed for each set of single organoids.

## Author Contributions

IC conceived of this project. IC analyzed project GENIE dataset. KL carried out all ddPCR experiments. MC generated organoids. KL, CS, KM, SM, and JP performed organoid isolations and digestion to single cells. All authors assisted in interpretation of data. KL, KS, and IC generated figures. KL and IC wrote manuscript with editorial support from all authors.

## Data Availability

Data supporting the findings of this study are available from the corresponding author upon reasonable request.

## Acknowledgements

This study was supported by funding provided by METAvivor, the Peter Carlson Trust, the Kristin L. Grampp Trust, Theresa’s Research Foundation, and the NCI/UTSW Simmons Cancer Center P30 CA142543. Special thanks to all members of the Chan Lab.

## Figure Legends

**Supplemental Figure 1: Invading and non-invading tumor organoids retain their morphology in liquid culture**

**A)** Schematic of organoid invasion assay for non-invading organoids (top) and invading organoids (bottom). Collagen was digested using collagenase for invading organoid conditions. Non-invading and invading organoids were pooled based on condition and gDNA isolated. **B)** Representative image of a non-invading organoid in liquid media. **C)** Representative image of an invading organoid in liquid media, post-collagenase digestion. Invasive organoids retain their invasive shape.

**Supplemental Figure 2: Copy number of ADGRA2 in tissue, pooled organoids, single cell digests, and single organoids from mouse mammary tumors**

**A)** ADGRA2 copy numbers in tissue, pooled organoids (Org), and single cell digests (Digest) from two different mice. Tissue sample copy number alterations are statistically different than pooled organoid samples (Mann-Whitney, p<0.05). **B)** ADGRA2 copy numbers in single organoids from tumors from two different mice.

**Supplemental Figure 3: Copy number of NSD3 in tissue, pooled organoids, single cell digests, and single organoids from mouse mammary tumors**

**A)** NSD3 copy numbers in tissue, pooled organoids (Org), and single cell digests (Digest) from two different mice. Tissue sample copy number alterations are statistically different than pooled organoid samples (Mann-Whitney, p<0.05). **B)** NSD3 copy numbers in single organoids from tumors from two different mice.

**Supplemental Figure 4: Copy number of PAK1 in tissue, pooled organoids, single cell digests, and single organoids from mouse mammary tumors**

**A)** PAK1 copy numbers in tissue, pooled organoids (Org), and single cell digests (Digest) from two different mice. No statistically significant differences in PAK1 copy number were appreciated (Mann-Whitney). **B)** PAK1 copy numbers in single organoids from tumors from two different mice.

