## Supplemental Figures for "Organoid generation from mouse mammary tumors captures the genetic heterogeneity of clinically relevant copy number alterations"

**Supplementary Figure 1**

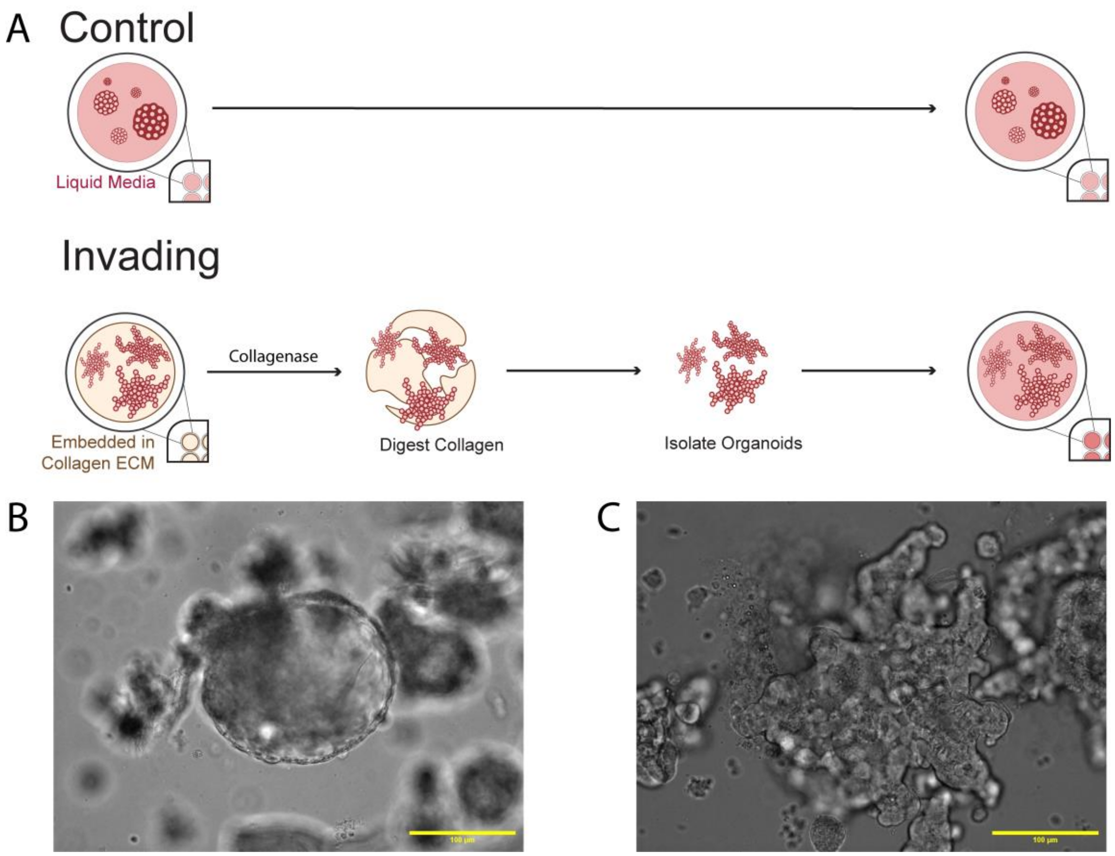


**Supplemental Figure 1: Invading and non-invading tumor organoids retain their morphology in liquid culture**

**A)** Schematic of organoid invasion assay for non-invading organoids (top) and invading organoids (bottom). Collagen was digested using collagenase for invading organoid conditions. Non-invading and invading organoids were pooled based on condition and gDNA isolated. **B)** Representative image of a non-invading organoid in liquid media. **C)** Representative image of an invading organoid in liquid media, post-collagenase digestion. Invasive organoids retain their invasive shape.

**Supplementary Figure 2**

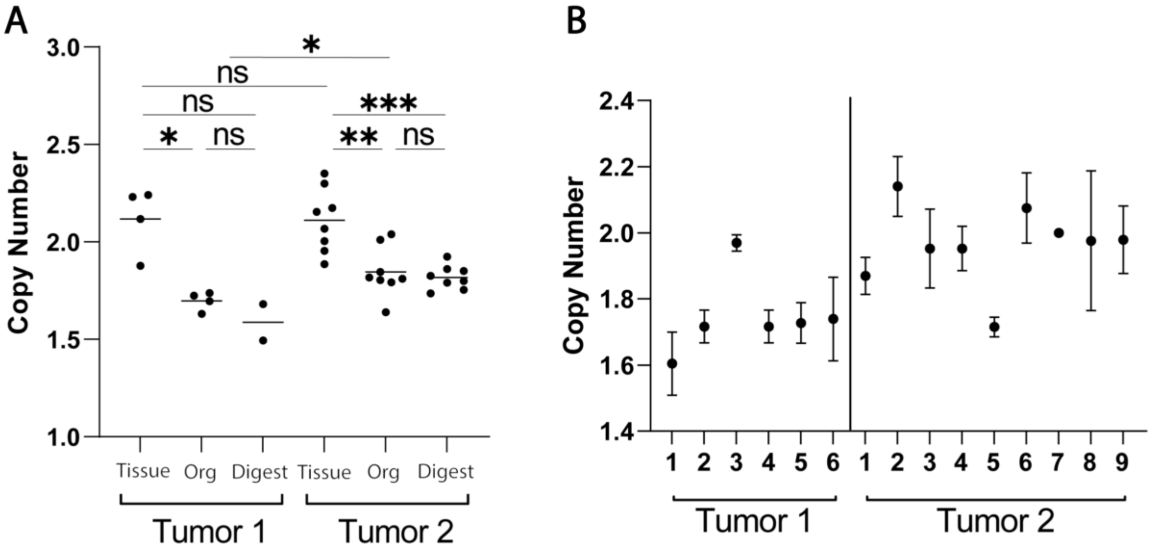


**Supplemental Figure 2: Copy number of ADGRA2 in tissue, pooled organoids, single cell digests, and single organoids from mouse mammary tumors**

**A)** ADGRA2 copy numbers in tissue, pooled organoids (Org), and single cell digests (Digest) from two different mice. Tissue sample copy number alterations are statistically different than pooled organoid samples (Mann-Whitney, $p<0.05$). **B)** ADGRA2 copy numbers in single organoids from tumors from two different mice.

**Supplementary Figure 3**

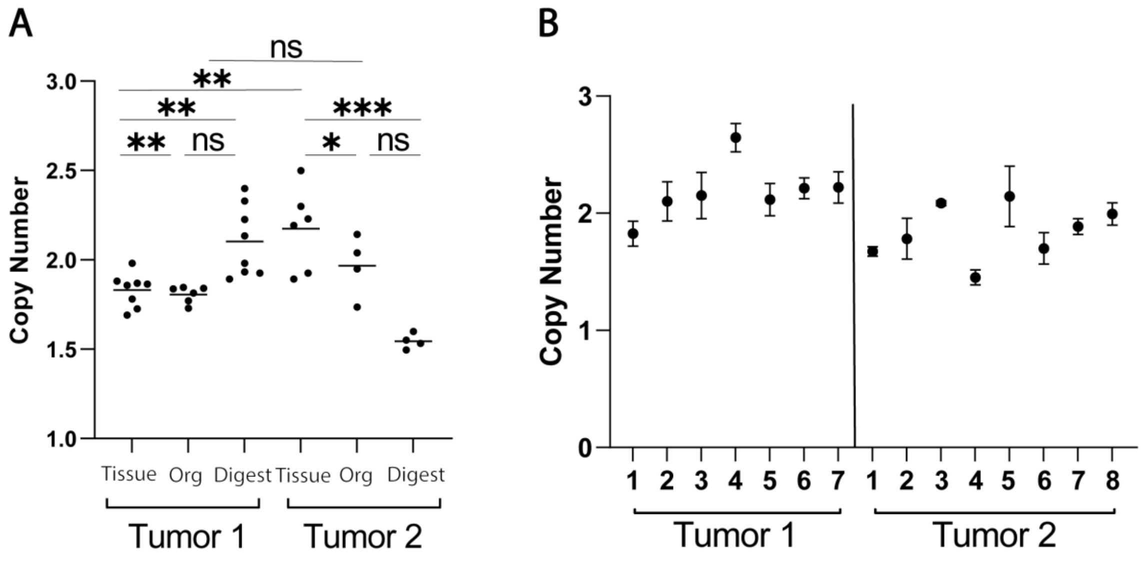


**Supplemental Figure 3: Copy number of NSD3 in tissue, pooled organoids, single cell digests, and single organoids from mouse mammary tumors**

**A)** NSD3 copy numbers in tissue, pooled organoids (Org), and single cell digests (Digest) from two different mice. Tissue sample copy number alterations are statistically different than pooled organoid samples (Mann-Whitney, $p<0.05$). **B)** NSD3 copy numbers in single organoids from tumors from two different mice.

**Supplementary Figure 4**

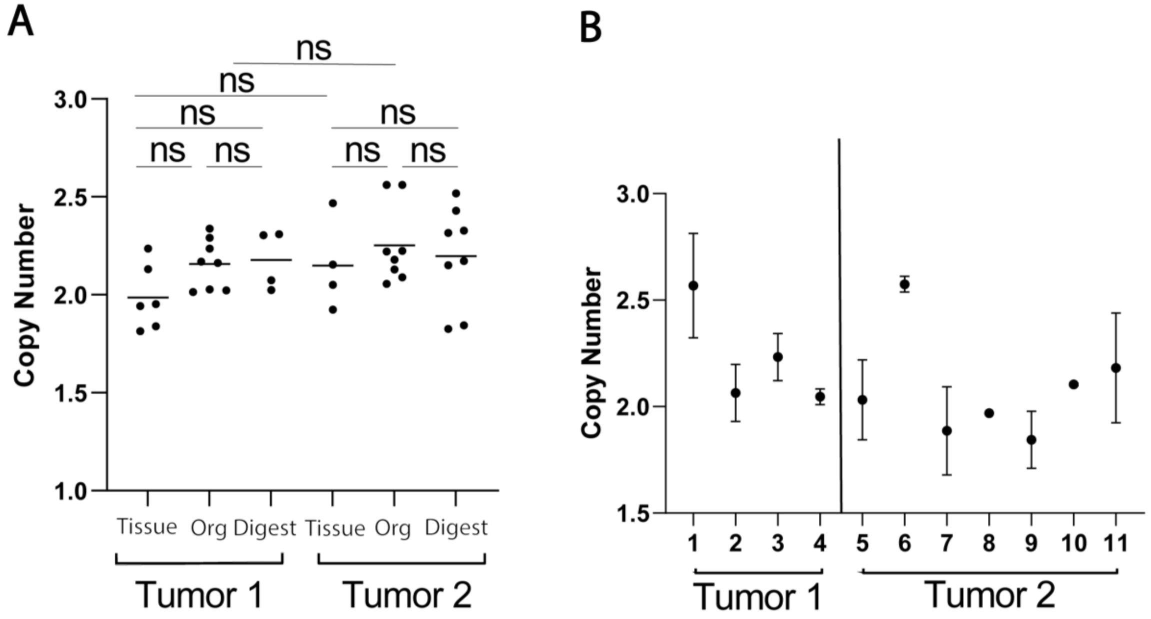


**Supplemental Figure 4: Copy number of PAK1 in tissue, pooled organoids, single cell digests, and single organoids from mouse mammary tumors**

**A)** PAK1 copy numbers in tissue, pooled organoids (Org), and single cell digests (Digest) from two different mice. No statistically significant differences in PAK1 copy number were appreciated (Mann-Whitney). **B)** PAK1 copy numbers in single organoids from tumors from two different mice.
